# PEBP regulates balance between apoptosis and autophagy, enabling coexistence of arbovirus and insect vector

**DOI:** 10.1101/2021.01.19.427364

**Authors:** Shifan Wang, Huijuan Guo, Keyan Zhu-Salzman, Feng Ge, Yucheng Sun

## Abstract

Apoptosis and autophagy are two most prominent forms of programmed cell deaths (PCD) that have been implicated in antiviral immunity in vertebrate and plant hosts. Arboviruses are able to coexist with its arthropod vectors by coordinating the PCD immunity, but the regulatory mechanism involved is largely unknown. We found that the coat protein (CP) of an insect-borne plant virus TYLCV directly interacted with a phosphatidylethanolamine-binding protein (PEBP) of its insect vector whitefly to negatively influence the MAPK signaling cascade. As a result, the apoptosis was activated in whitefly which increased viral loading. Simultaneously, the PEBP4-CP interaction liberated ATG8, the hallmark of autophagy initiation, and eliminates arbovirus. Furthermore, apoptosis-promoted virus loading was compromised by agonist-induced autophagy, but autophagy-associated suppression on virus loading was unaffected by apoptosis agonist or inhibitor, suggesting that virus loading was predominantly determined by autophagy rather than by apoptosis. Our results demonstrated that maintaining a mild immune response by coordinating apoptosis and autophagy processes presumably could facilitate coexistence of the arbovirus and its insect vector. Taken together, immune homeostasis shaped by two types of PCD may facilitate the arbovirus preservation within the insect vector.

**Graphical abstract:** 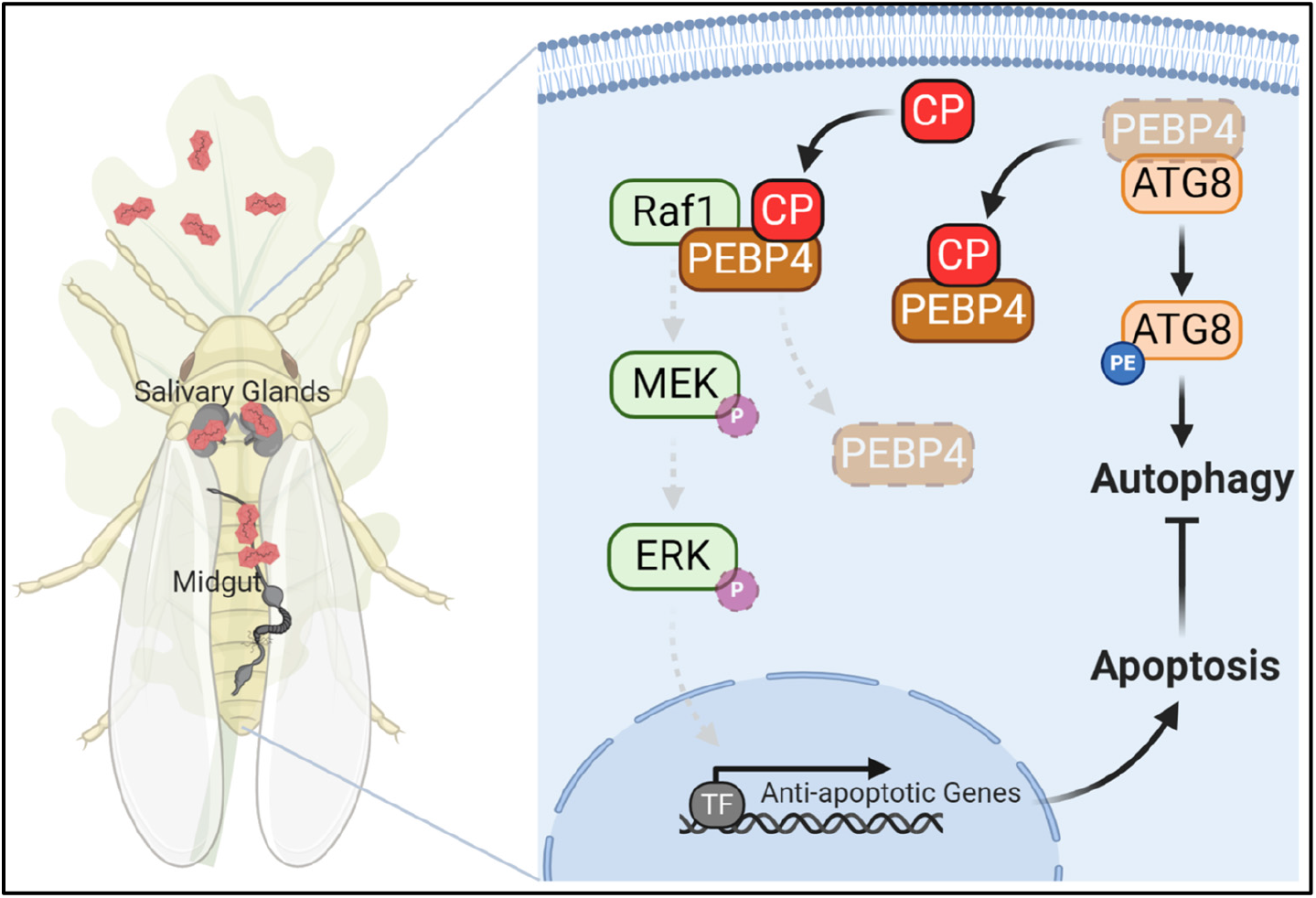

**Highlights:**

- Interaction between whitefly PEBP4 and TYLCV CP suppresses phosphorylation of MAPK cascade, activating apoptosis
- TYLCV CP liberates PEBP4-bound ATG8, resulting in lipidation of ATG8 and initiation of autophagy.
- PEBP4 balances apoptosis and autophagy in viruliferous whitefly to optimize virus loading without obvious fitness cost.

## Introduction

Programmed cell death (PCD), including apoptotic and autophagic cell death, is an essential process of innate and adaptive immunity, protecting vertebrates from pathogen infection (Deretic and Levine, 2009; Fuchs and Steller, 2011; Jorgensen et al., 2017; Lockshin, 2016; Nagata and Tanaka, 2017; Virgin and Levine, 2009; Zhou and Zhang, 2012). Autophagy involves sequestration of portions of the cytoplasm by autophagosomes, the double-membrane vesicles, which fuse with lysosomes to form autolysosomes in which autophagic cargo is degraded (Mizushima and Komatsu, 2011; Yang and Klionsky, 2010). This lysosomal degradation mechanism contributes to cell differentiation, development, and homeostasis(Taylor et al., 2008). Apoptosis is the most extensively studied PCD, it has been characterized by cytoplasmic shrinking, cell rounding, chromatin condensation, DNA fragmentation and membrane blebbing (Fuchs and Steller, 2011; Jorgensen et al., 2017; Lockshin, 2016). The major types of apoptosis are orchestrated by cysteine proteases caspases, and respectively triggered by mitochondria-dependent intrinsic and death receptor-dependent extrinsic pathways (Bock and Tait, 2020; Taylor et al., 2008). Initiation of apoptosis via the effector caspases activates catabolic hydrolases that can degrade most of the macromolecules of the cell (Galluzzi et al., 2009). In most cases, PCD-associated degradation serves as an effective barrier to ward off invading pathogens but usually causes certain tissue damages in hosts (Griffith and Ferguson, 2011). By contrast, PCD in insect vectors seems more compatible with arthropod-borne virus (arbovirus) with less or even no pathological symptoms during virus acquisition and transmission(Bartholomay and Michel, 2018; Brackney, 2017; Clem, 2016; Yang et al., 2020). Evidence suggests that arbovirus-induced PCD in insect vectors enhances the immune tolerance, while permitting coexistence of arbovirus with the vector (Chen et al., 2019; Chen et al., 2017; Wang et al., 2020c). The molecular and biochemical bases, however, remain largely elusive.

Controlled cell destruction is often considered as an innate defense mechanism that eliminates the need for infected cells to counteract viral infection. Therefore, many viruses suppress apoptosis to prevent premature host cell death and thus enhance virus proliferation (Lockshin, 2016). By contrast, the activation of apoptosis in insect vectors is usually thought to be beneficial to arbovirus. For instance, suppression of apoptosis in *Aedes aegypti* decreases the virus loading and impairs dissemination of dengue virus (Eng et al., 2016). Activation of apoptosis in midgut could damage the integrity of gut barrier of mosquito and modify the cellular regeneration program, leading to establishment of an accessible route to the vector’s hemolymph (Janeh et al., 2017; Taracena et al., 2018). Furthermore, crosstalk between apoptosis and autophagy also regulates virus invasion in infected hosts (Maiuri et al., 2007; Marino et al., 2014). Although autophagic cell death (or type II cell death) is an alternative pathway to cellular demise, autophagy itself normally suppresses apoptosis to constitute a stress adaption that avoiding cell death (Maiuri et al., 2007; Marino et al., 2014). In contrast to apoptosis, arbovirus-induced autophagy could function as intracellular resistance mechanism to eliminate arbovirus infection and replication. It is exemplified by tomato yellow leaf curl virus (TYLCV) that is primarily transmitted by whitefly (Wang et al., 2016). TYLCV activates autophagy in the midgut and salivary glands of the Middle East Asia Minor 1 (MEAM1) whitefly species, and suppression of autophagy increases the TYLCV abundance (Wang et al., 2016). In these cases, apoptosis and autophagy can be separately stimulated by arbovirus invasion. Considering the complexity of interplay between these two forms of PCD, it conceives that apoptosis and autophagy could occur simultaneously and are orchestrated in spatial and temporal *in vivo*, which could be a crucial regulatory basis for maintaining a balance of vector and virus loading.

Antagonism between apoptosis and autophagy has been reported in most cases despite other scenarios are also shown, i.e., apoptosis preceding autophagy, autophagy activating apoptosis (Maiuri et al., 2007; Marino et al., 2014). A number of crosstalk points have been identified including Beclin-1 and Bcl-2/Bcl-xL, FADD and Atg5, caspase- and calpain-mediated cleavage of autophagy-related proteins, and autophagic degradation of caspases (Gordy and He, 2012). Among them, Bcl-2 family that functions in maintaining mitochondrial outer membrane permeabilization (MOMP) and endoplasmic reticulum (ER) calcium stores serves as pivotal regulatory molecules for homeostatic balance of PCD (Bock and Tait, 2020; Chipuk et al., 2010; Gordy and He, 2012). This family comprises proapoptotic/effector proteins (Bax and Bak), antiapoptotic proteins (Bcl-2, Bcl-xL, Mcl1, Bcl-W and A1), and BH3-only proteins (Bim, Bad, Bmf, Bid and PUMA) (Chipuk et al., 2010; Lavoie et al., 2020). The ratio of these molecules could be regulated by several signaling pathways, such as MAPK (Raf1/MEK/ERK), which finally controls the homeostatic balance of apoptosis and autophagy and regulates the process of PCD (Chipuk et al., 2010; Lavoie et al., 2020). For instance, ERK typically inhibits cell death via intrinsic pathway of apoptosis in ways that suppresses proapoptotic proteins or stimulates antiapoptotic proteins (Lavoie et al., 2020).

The whitefly (*Bemisia tabaci*) is a notorious insect vector worldwide and transmits multiple DNA viruses belonging to Geminiviridae, the most devastating pathogens infecting hundreds of crops (Rosen et al., 2015). The MEAM1 and Mediterranean (MED) whitefly species transmitted TYLCV a monopartite begomovirus (family Geminiviridae) in a persistent-propagative manner (Rosen et al., 2015). Previous results showed that both apoptosis and autophagy could be activated by TYLCV in MEAM1 whitefly (Wang et al., 2016; Wang et al., 2020b). While virus loading is facilitated by activation of apoptosis, it is apparently suppressed by initiation of autophagy (Wang et al., 2016; Wang et al., 2020a; Wang et al., 2020c). The underlying regulatory mechanism and ecological implication, however, remain largely unclear. Here, we showed that once bound with TYLCV CP, the phosphatidylethanolamine-binding protein 4 (PEBP4) of whitefly interacted with Raf1 to suppress the phosphorylation of MAPK signaling and thereby activated apoptosis. Meanwhile, binding to CP caused dissociation of PEBP4 from ATG8, which triggered the autophagy pathway. Simultaneously induction of apoptosis and autophagy by TYLCV in whitefly optimized arbovirus abundance such that maximal fitness of the vector could be achieved. Our results revealed a novel function of PEBP4 in regulating whitefly immune homeostasis.

## Results

### TYLCV simultaneously induced apoptosis and autophagy in whitefly

Since the midgut and the salivary gland were important barriers for arbovirus circulation within the insect vector, they were dissected to determine the induction dynamics of apoptosis and autophagy in viruliferous whitefly. The TUNEL assay and immunofluorescence results showed that both apoptosis and autophagy were activated in the midgut at 24 hours post-infection (hpi), and were enhanced in salivary gland which possessed a basal fluorescence intensity of apoptosis and autophagy (Figure 1A and Figure 1B). Viral acquisition decreased the expression of anti-apoptotic genes *Iap* and *Bcl-2*, but increased the expression of pro-apoptotic genes *Caspase1* and *Caspase3* (Figure 1C). Furthermore, the autophagy-related genes *ATG3*, *ATG8*, *ATG9*, and *ATG12* were also up-regulated in viruliferous whitefly (Figure 1D). Consistently, activation of apoptosis and autophagy at the protein level were further confirmed by the time-course immunoblotting assay. Typically, the full-length Caspase3 was rapidly converted to cleaved Caspase3 as the time increases from 6 to 24 hpi, suggesting a quickly induced apoptosis in viruliferous whitefly since 6 hpi. Likewise, a remarkable consumption of autophagy substrate SQSTM1 and increase of ATG8 lipidation (conjugated to the phosphatidylethanolamine, ATG8-PE) were also observed from 6 to 24 hpi, suggesting a simultaneous induction of autophagy, along with the apoptosis (Figure 1E).

**Figure 1.**
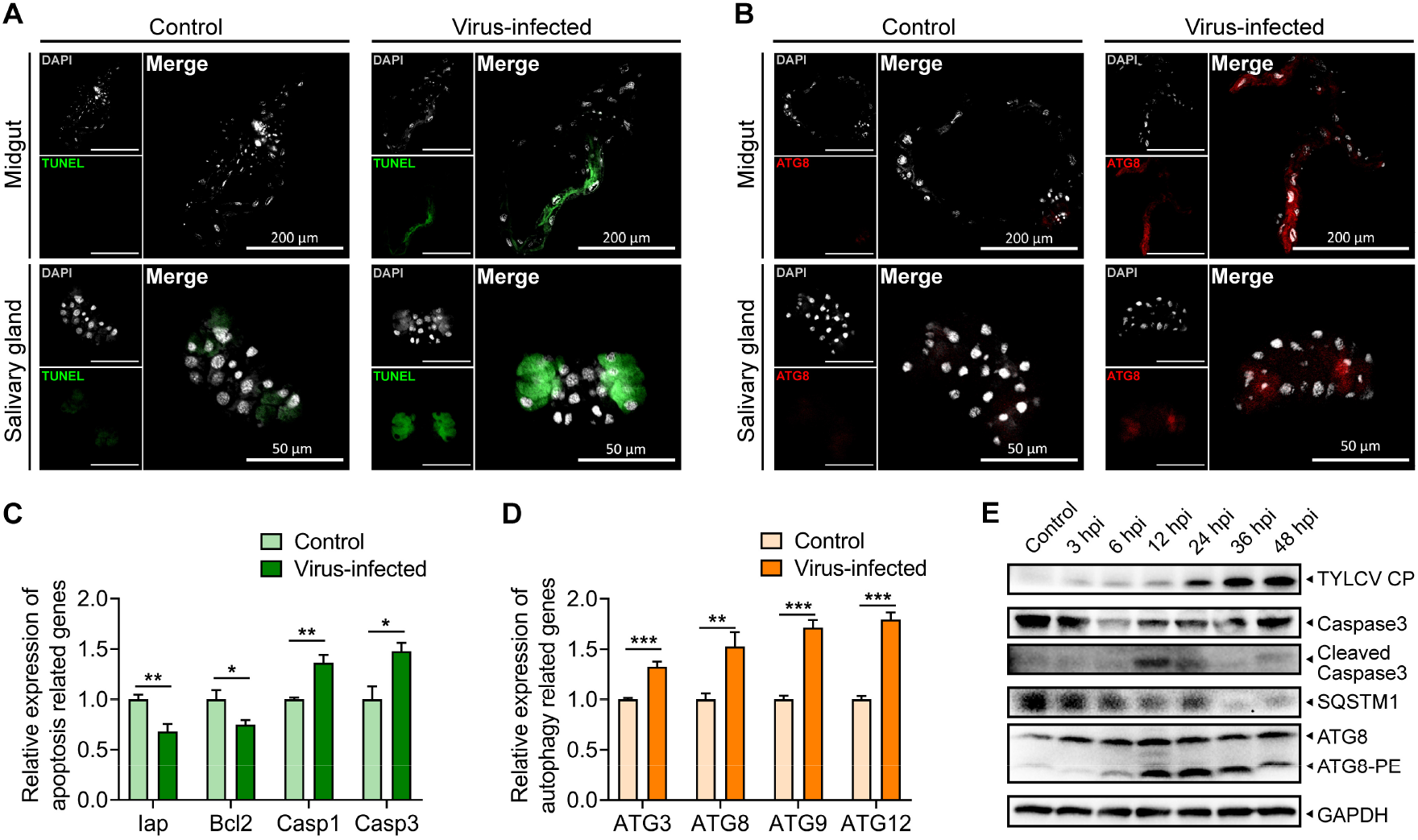
Apoptosis and autophagy were simultaneously induced in whitefly when acquiring TYLCV. Midguts and salivary glands were dissected from nonviruliferous or viruliferous whitefly. (A) Apoptosis was determined by TUNEL assay (green), and (B) autophagy was labelled by the hallmark ATG8 (red). The nuclei (white) were stained by DAPI. n≥8. Whiteflies fed with TYLCV-infected plant for 24 hours were sampled. The relative expressions of (C) anti-apoptotic genes (*Iap*, *Bcl2*), pro-apoptotic genes (*Caspase1*, *Caspase3*), and (D) autophagy related genes (*ATG3*, *ATG8*, *ATG9*, *ATG12*) were determined by qPCR. Values in bar plots represent mean ± SEM (\**P*<0.05, \*\**P*<0.01, \*\*\**P*<0.001), n=5. (E) Time-course immunoblot monitored the dynamics of apoptosis and autophagy activation, and GAPDH served as the loading control. The consumption of SQSTM1, a crucial autophagy substrate, represented autophagy activation. ATG8-PE represented the lipidation of ATG8, the hallmark of autophagy, in forms of ATG8 combining with phosphatidylethanolamine (PE).

### PEBP4 directly interacted with the coat protein (CP) of TYLCV

CP has been shown to be required for successful infection (Hogenhout et al., 2008), it therefore is used as the bait in yeast two-hybrid (Y2H) to screen the target proteins in whitefly that may potentially be responsible for the activation of apoptosis and autophagy. Twenty-three nonredundant sequences were firstly blasted against the genome of MED whitefly (http://gigadb.org/dataset/100286) to acquire the full amino acid sequences, which were then annotated referring to the database of Hemiptera in NCBI (Table S1). Of all the potential proteins with high sequence identities, we focused on a putative phosphatidylethanolamine-binding protein (PEBP) due to its putative function in PCD regulation (Noh et al., 2016; Role, 2004; Schoentgen and Jonic, 2018). The phylogenetic analysis showed that this PEBP was homologous to the PEBP4 family with a conserved PE-binding domain, and hereafter was named as whitefly PEBP4 (Figure S1A and Figure S1B).

Interaction between TYLCV CP and PEBP4 was further verified using Y2H analysis. Plating on triple dropout (TDO) medium, growth was observed in colonies that harbored both proteins but not for high-stringency condition of quadruple dropout (QDO) medium (Figure 2A). Two polyclonal PEBP4 custom antibodies against different synthetic peptides were used in the co-immunoprecipitation (CoIP) assay to corroborate the virus CP and whitefly PEBP4 interaction *in vivo*. The CoIP results showed that TYLCV CP was co-immunoprecipitated with PEPB4 in viruliferous whitefly lysate rather than non-viruliferous whitefly lysate, regardless of which PEPB4 antibody was used (Figure 2B). To determine the domain-specific interaction with CP, whitefly PEBP4 was expressed as two protein peptides: a species-specific pre-domain and a conserve PE-binding domain. GST pull-down assay found that the pre-domain of PEBP4, rather than the PE-binding domain, interacted with the virus CP, suggesting this protein-protein interaction may be MED whitefly-specific (Figure 2C). Immunofluorescence showed that PEBP4 and virus CP co-localized in midgut and salivary gland (Figure 2D). These results further confirmed direct CP-PEBP4 interaction.

**Figure 2.**
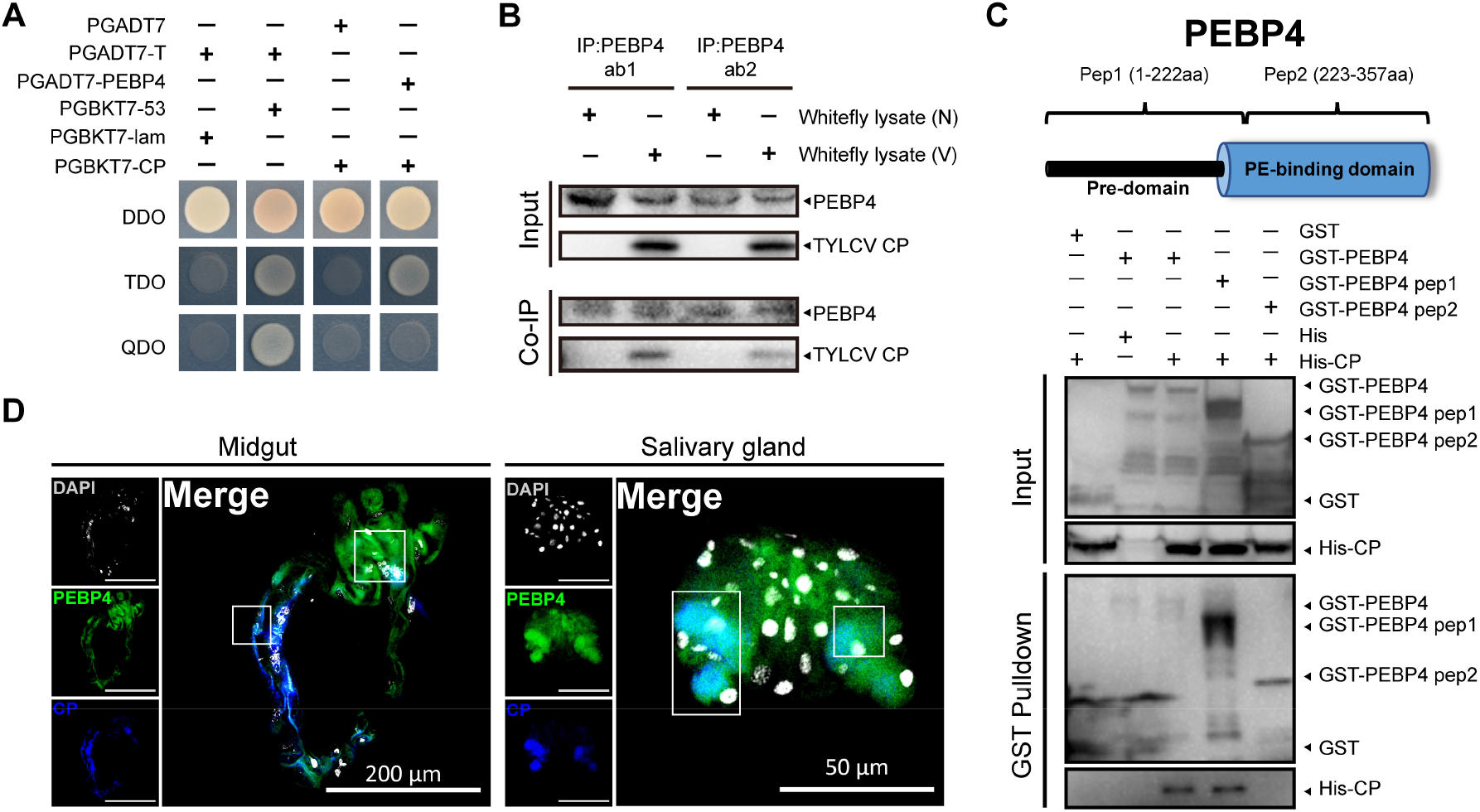
PEBP4 directly interacted with the CP of TYLCV. (A) PEBP4 interacted with TYLCV CP in yeast two-hybrid assay. Lane 1: negative control, lane 2: positive control, lane 3: blank group, lane 4: PEBP4-CP interaction group. (B) CoIP assay. Total proteins of the viruliferous (V) vs. non-viruliferous (N) whitefly were extracted. Two polyclone antibodies (ab1, ab2) of whitefly PEPB4 were used. (C) GST pull-down assay. Full length PEBP4 was divided and expressed separately as pep1 and pep2. The pep1 consisted of a whitefly specific pre-domain while pep2 had a conserved PE-binding domain. Recombinant His-CP interacted with both GST-PEBP4 and GST-PEPB4 pep1, instead of GST-PEPB4 pep2. The GST tag and His tag expressed by empty vector were used as negative control. (D) PEPB4 (green) was typically expressed in the midgut and salivary gland of whitefly, in which PEPB4 was co-localized with the CP of TYLCV (blue). The nuclei (white) were stained by DAPI, n≥7.

### PEBP4 knockdown in viruliferous whitefly attenuated TYLCV-induced apoptosis but enhanced autophagy

The abundance of PEBP4 decreased at the very beginning of TYLCV acquisition while increased from 12 to 24 hpi at both transcriptional and translational levels (Figure S2A and Figure S2B). Feeding on *dsPEBP4* decreased *BtPEBP4* expression in whitefly (Figure 3A). Knockdown of *BtPEBP4* increased gene expression of anti-apoptotic *Iap* and *Bcl2* while reduced pro-apoptotic *Caspase1* and *Caspase3*, indicating that apoptosis in whitefly was positively regulated by PEBP4 (Figure 3B). By contrast, autophagy-related genes *ATG3* and *ATG12* were increased when fed with *dsPEBP4* relative to *dsGFP*, indicating autophagy was negatively regulated by PEBP4 (Figure 3C). Fluorescence data showed that the virus-induced apoptosis was suppressed but autophagy increased in both midgut and salivary gland (Figure 3D, 3E). The immunoblotting results fully agreed with those of immunofluorescence assays in that knockdown of *BtPEBP4* resulted in increased Caspase3 cleavage, SQSMT1 consumption and ATG8 lipidation, as well as decreased TYLCV loading in viruliferous whitefly (Figure 3F). Taken together, apoptosis and autophagy in whitefly could be independently regulated by PEBP4-CP interaction.

**Figure 3.**
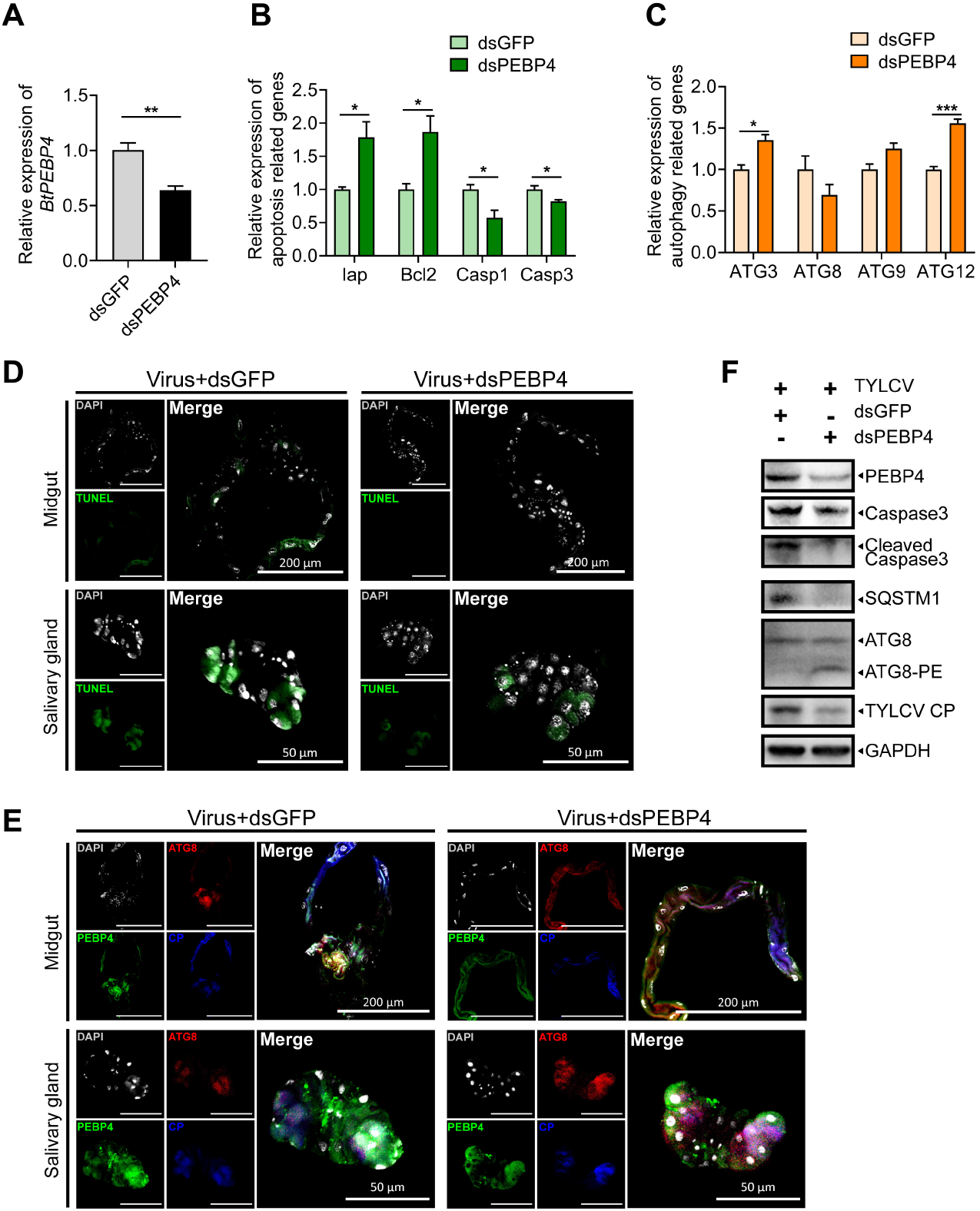
Knockdown of PEBP4 attenuated TYLCV-induced apoptosis but enhanced autophagy. (A) *BtPEBP4* RNAi knockdown in whitefly reduced the expression of *BtPEBP4* at mRNA level, and *dsGFP* served as control. (B) The relative expression of apoptotic related genes (*Iap*, *Bcl2*, *Caspase1*, *Caspase3*), and (C) autophagy related genes (*ATG3*, *ATG8*, *ATG9*, *ATG12*) were determined using qPCR. Values in bar plots represent mean ± SEM (\**P*<0.05, \*\**P*<0.01, \*\*\**P*<0.001), n=3. (D) Apoptosis in midgut and salivary gland were determined by TUNEL labeling (green). (E) Autophagy was determined by immunofluorescence: PEPB4 (green), ATG8 (red), TYLCV CP (blue), DAPI (white), n≥8. (F) Apoptosis and autophagy in whole body of whitefly were determined by immunoblotting. GAPDH served as loading control, and three independent biological replicates were conducted.

### PEBP4 interacted with Raf1 and negatively regulated MAPK pathway phosphorylation to enhance the virus-induced apoptosis

The MAPK phosphorylation pathway Raf1/MEK/ERK relays the regulation of PEBPs on apoptosis initiation (Role, 2004; Schoentgen and Jonic, 2018). All members of the human PEBP family are capable of regulating the MAPK cascade (Dai et al., 2016; Hickox et al., 2002a; Hu et al., 2014; Su et al., 2016; Zuo et al., 2015). To investigate whether whitefly PEBP4 regulated the TYLCV-induced apoptosis via this signaling pathway, we examined if there existed direct interaction between PEBP4 and Raf1 *in vivo* and *in vitro*. GST pull-down and CoIP assays showed that PEBP4 interacted with Raf1 via its conserve PE-binding domain regardless of CP existence, suggesting the conserved PE-binding domain was required for this interaction with Raf1 (Figure 4A and Figure 4B). TYLCV infection increased expression of *Raf1* and *MEK* but was not significant for *ERK* (Figure S3A). By conducting a time-course virus acquisition experiment, the immunoblot results showed that TYLCV suppressed the phosphorylation of ERK and MEK (Figure 4C). Knockdown of PEBP4 enhanced the phosphorylation, indicating that PEBP4 was a negative regulator of this signaling pathway (Figure S3B). Since TYLCV could increase the transcripts of unphosphorylated Raf1 (Figure S3A), it is uncertain that the suppression on MAPK phosphorylation at protein level still was resulted from PEBP4-Raf1 interaction. Whiteflies therefore were lysed and co-incubated with a concentration gradient of GST-PEBP4 and His-CP respectively to rule out the transcriptional *do novo* synthesis of unphosphorylated Raf1. The results showed that both PEBP4 and TYLCV CP directly increased the unphosphorylated Raf1, which equals to decrease phosphorylated Raf1 because no additional Raf1 products could be supplied in whitefly lysate, resulting in reducing the phosphorylation of Raf1 and negatively regulating MAPK signaling cascade (Figure 4D and Figure 4E). Taken together, TYLCV infection not only increased unphosphorylated Raf1at transcriptional level but directly reduced the phosphorylation of Raf1 at protein level. The CoIP results showed that neither CP nor Raf1 negatively affected the interaction with PEBP4, indicating the competitively bound with PEBP4 was not shown for CP and Raf1 in viruliferous whitefly (Figure 4F). Similarly, in inflorescence results, PEBP4 and Raf1 merged in midgut and salivary gland regardless TYLCV CP existence or absence, and CP could merge with PEPBP4 and Raf1, suggesting a triple complex CP-PEBP4-Raf1 could exist (Figure 4G).

**Figure 4.**
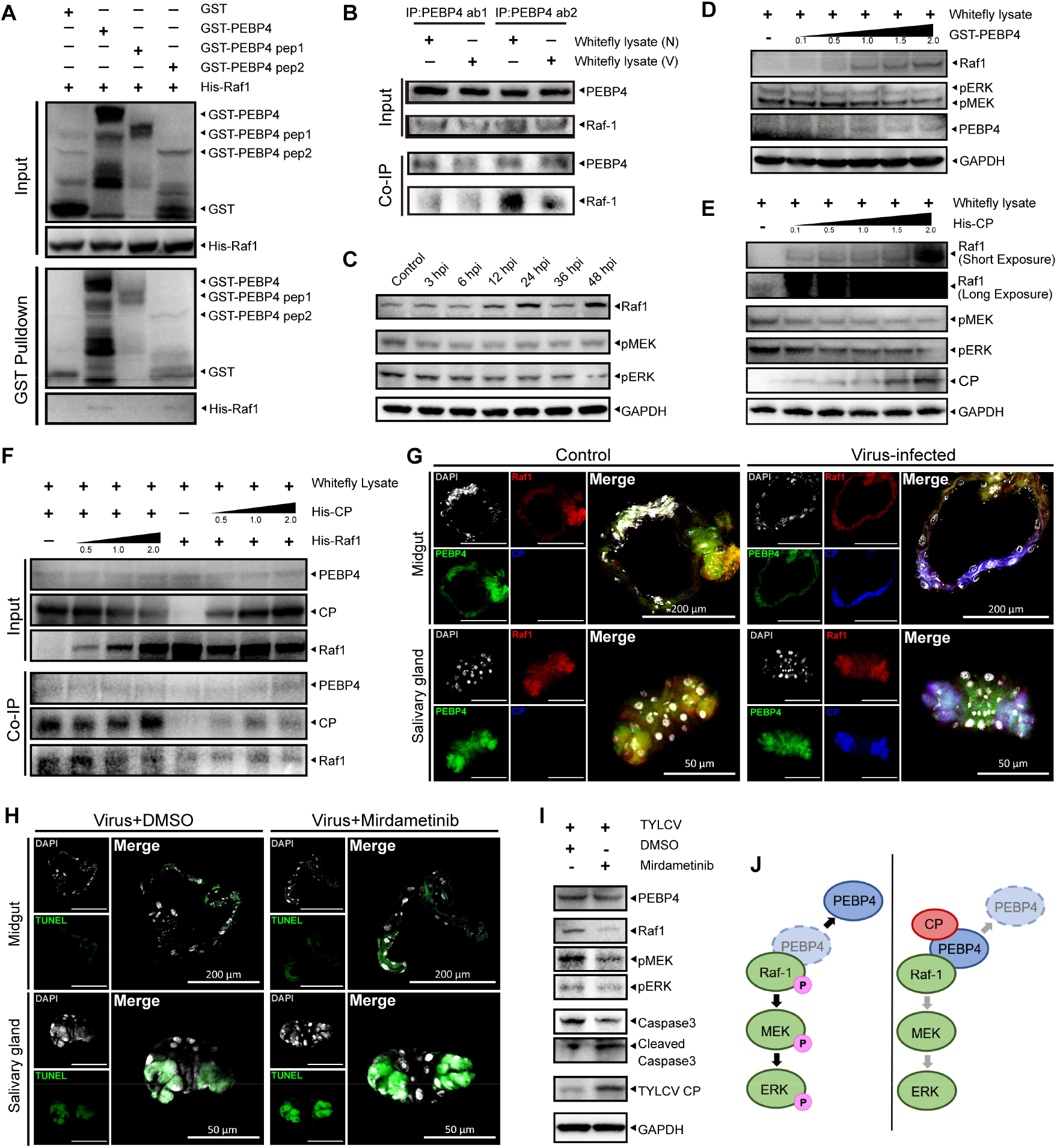
PEBP4 interacted with Raf1 which negatively regulated MAPK phosphorylation and enhanced the virus-induced apoptosis. (A) GST pull-down showing the interaction between recombinant His-Raf1 and PE-binding domain of GST-PEBP4. (B) PEBP4 co-immunoprecipitated with Raf1 in viruliferous (V) or nonviruliferous (N) whitefly. (C) Time-course immunoblots of MAPK pathway phosphorylation. (D-E) Immunoblots of MAPK pathway phosphorylation in whitefly lysate when co-incubated with a concentration gradient of (D) PEBP4 and (E) TYLCV CP. (F) His-CP and His-Raf1 with different concentrations were co-incubated in whitefly lysate in CoIP assay. (G) Raf1 (red), PEBP4 (green), and CP (blue) were simultaneously labeled in immunofluorescence assay, n≥8. (H-I) After 24 h feeing on mirdametinib, a phosphorylation inhibitor of MEK, whiteflies were transferred to virus-infected host plants for another 24 hours for TYLCV acquisition. The midgut and salivary gland were dissected for (H) TUNEL labeling (green). The nuclei were stained by DAPI (white), n≥6. (I) Immunoblot of phosphorylation of MAPK pathway and initiation of apoptosis. (J) Model of CP-PEBP4 regulation on MAPK cascade in whitefly. CP stabilized Raf1 interaction with PEBP4, which disabled the MAPK phosphorylation cascade.

MAPK pathway has been shown to negatively regulate apoptosis in most cases (Kale et al., 2018; Maiuri et al., 2007; Marino et al., 2014). ERK-regulated anti-apoptotic genes, including *Iap* and *Bcl-2*, normally forming a balance with pro-apoptotic genes to prevent the apoptosis (Bock and Tait, 2020; Taylor et al., 2008). Suppression of anti-apoptotic genes could increase the ratio of pro-apoptotic genes products and eventually induced apoptosis (Kale et al., 2018; Lavoie et al., 2020; Maiuri et al., 2007; Marino et al., 2014). We hypothesized that phosphorylation of MAPK pathway negatively regulated the TYLCV-induced apoptosis, and conversely, suppression of MAPK pathway phosphorylation in whitefly could promote the arbovirus-induced apoptosis. The immunofluorescence and immunoblot results showed that TYLCV-induced apoptosis was enhanced when the phosphorylation of MAPK signaling cascade was suppressed by Mirdametinib, an inhibitor of MEK phosphorylation (Figure 4H and 4I). These results revealed that, the presence of TYLCV CP stabilized PEBP4 binding to Raf1 via its PE-binding domain, which suppressed the phosphorylation of MAPK signaling cascade and promoted apoptosis (Figure 4J).

### TYLCV CP competitively interacted with PEBP4 to liberate ATG8 and promoted autophagy

On the resting state, human PEBP1 interacts with LC3B, the microtubule associated protein 1 light chain 3 β, at the LC3-interacting region (LIR) motif to suppress autophagy in a MAPK pathway-independent manner (Noh et al., 2016). Once phosphorylated at Ser153, PEBP1 dissociates from the PEBP1-LC3B complex and induces autophagy (Noh et al., 2016). Here, we found that whitefly PEBP4 also contained putative LIR motifs based on the prediction of iLIR Autophagy Database (https://ilir.warwick.ac.uk/index.php) (Figure S4). The CoIP and GST-pulldown assays demonstrated that whitefly PEBP4 interacted with ATG8, the LC3 homolog in insects, both *in vivo* and *in vitro* (Figure 5A, 5B). The *in vitro* co-incubation assays showed enhanced autophagy as recombinant CP increased (Figure 5C). By contrast, whitefly lysate co-incubated with PEBP4 had little effect on autophagy without TYLCV induction, suggesting that PEBP4 in nonviruliferous whitefly was occupied by ATG8 on resting state leading to inactivated autophagy (Figure 5D). When activated autophagy by recombinant CP, CP interacted with PEBP4 and released ATG8, and excess supplement of PEBP4 can rescue ATG8 liberation, and thereby suppressed virus-induced autophagy, suggesting contrasting effects of CP and PEBP4 on autophagy activation (Figure 5E). It is reasonable to speculate that virus CP could competitively interact with PEBP4 to dissociate PEBP4 and ATG8 interaction. A competitive pull-down assay was therefore conducted. ATG8 overdose had little effect on PEBP4-CP interaction while excess of virus CP impaired the PEBP4-ATG8 association (Figure 5F). It appeared that PEBP4 had higher affinity to TYLCV CP than to ATG8. Similar results were corroborated by immunofluorescence evidence that activated ATG8 was increased in midgut and salivary gland once acquiring TYLCV (Figure 5G). These findings suggested that autophagy was inactivated in nonviruliferous whitefly because ATG8 was arrested by PEBP4. Once infected by TYLCV, the virus CP competitively interacted with PEBP4 and liberated ATG8 that supposedly underwent lipidation and initiated the formation of autophagosome (Figure 5H).

**Figure 5.**
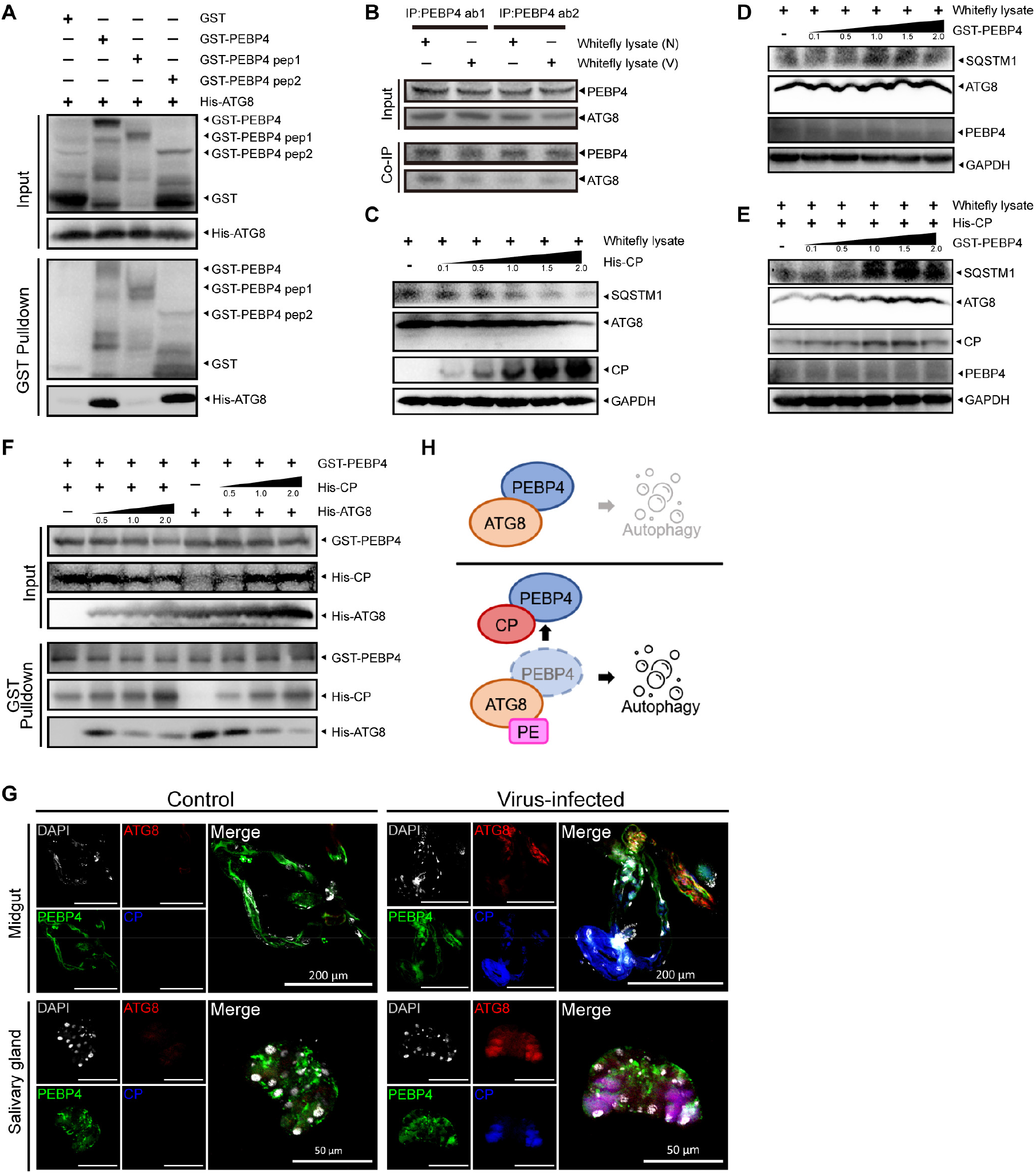
CP competitively interacted with PEBP4 to liberate ATG8 and promoted autophagy. (A) GST pull-down showing the interaction between recombinant His-ATG8 and GST-PEBP4. (B) PEBP4 co-immunoprecipitated with ATG8 in viruliferous (V) or nonviruliferous (N) whitefly. (C-E) Recombinant His-CP and GST-PEBP4 with a concentration gradient were co-incubated with the lysate of whitefly to monitor the activation of autophagy by immunoblotting. (F) The competition between CP and ATG8 on PEPB4 interaction was detected by a competitive pull-down assay. His-CP and His-ATG8 were supplied within a concentration gradient to co-incubate with GST-PEBP4. (G) ATG8 (red), PEBP4 (green), and CP (blue) were simultaneously labeled and localized by immunofluorescence in viruliferous whitefly. The nuclei were stained by DAPI (white), n≥8. (H) Model of CP-PEBP4 regulation on ATG8 in whitefly. PEBP4 arrested with PE-unconjugated ATG8 to prevent autophagy in nonviruliferous whitefly. Once TYLCV invasion, CP disassociated with PEBP4-bound ATG8, which led to the liberation of ATG8 and in turn initiation of autophagy.

### A mild PCD response ensured coexistence of TYLCV and whitefly

It has been shown that apoptosis enhanced TYLCV loading while autophagy eliminated in MEAM1 whitefly (Wang et al., 2016; Wang et al., 2020c), but is still unclear whether simultaneous activation of apoptosis and autophagy was essential in MED whitefly for co-existence with TYLCV. We therefore introduced a serial of pharmacological experiments to artificially regulate the apoptosis and autophagy of whitefly, and to determine effects of these PCD on virus loading. The results of immunoblots and qPCR showed that autophagy inhibitor 3-MA and apoptosis agonist PCA-1 respectively increased the abundance of CP and genomic DNA of TYLCV in viruliferous whitefly. By contrast, autophagy agonist rapamycin and apoptosis inhibitor Z-VAD-FMK respectively reduced the relative abundance of CP and genomic DNA of TYLCV (Figure 6A and Figure 6B). These results confirmed that attenuation of autophagy and/or activation of apoptosis increased TYLCV abundance in whitefly.

**Figure 6.**
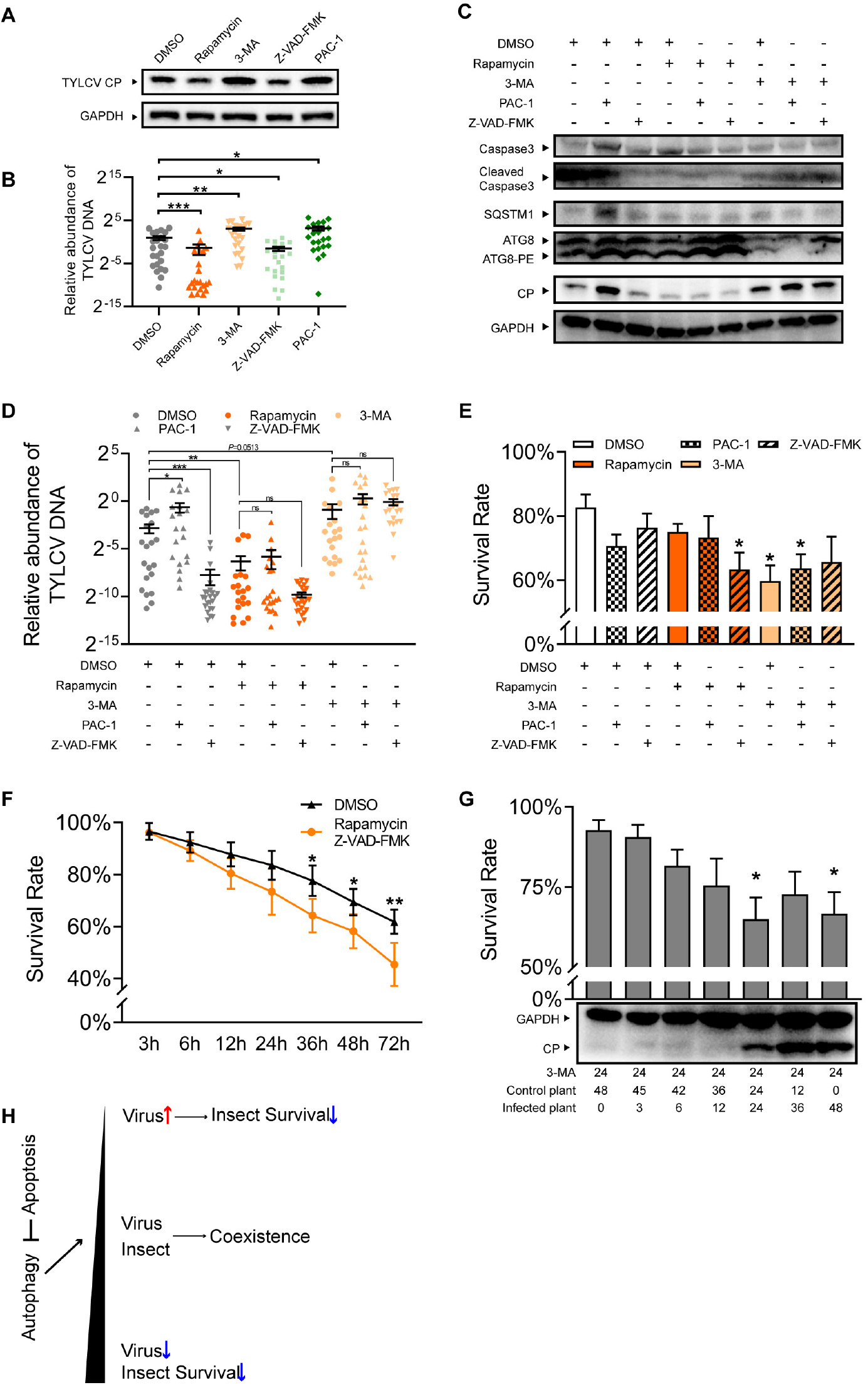
A mild intracellular immunity in whitefly facilitated its coexistence with TYLCV. (A-B) Autophagy agonist rapamycin, autophagy inhibitor 3-MA, apoptosis agonist PAC-1, and apoptosis inhibitor Z-VAD-FMK were fed with 24h in artificial diet respectively. (A) The relative protein abundance of TYLCV CP was determined by immunoblotting in viruliferous whitefly, and (B) the relative quantity of TYLCV DNA per whitefly was determined by qPCR, n=21-24. (C-E) Effects of pharmacological combinations of agonist and inhibitor on activation of apoptosis and autophagy in viruliferous whitefly, and their consequences to virus loading and survival rate of whitefly. Whiteflies were transferred to acquire virus for 24 h after feeding with agonist or/and inhibitor. (C) The activation of apoptosis and autophagy in viruliferous whitefly were determined by immunoblotting, and (D) the relative quantity of virus DNA in whitefly was determined by qPCR, n=21-24. (E) 100 whiteflies per replicate from each pharmacological combination treatment were transferred to virus-infected plant for 48 h, and then was sampled for survival rate measurement, n=3. (F) Nonviruliferous whiteflies fed with rapamycin and Z-VAD-FMK in combination were sampled for survival rate measurement in terms of over-activation of autophagy, n=3. (G) To determine whether overloading of TYLCV could directly impair survival rate of whitefly, autophagy inhibitor 3-MA was fed with 24 h. Each group consisting of 60 whiteflies was transferred to uninfected plant, and then was transferred again to the TYLCV-infected plant following the time point set. The survivals were recorded for survival rate calculation, and were sampled for determination of TYLCV CP abundance by immunoblotting, n=3. Values in all bar or dot plots represent mean ± SEM (\**P*<0.05, \*\**P*<0.01, \*\*\**P*<0.001). (H) Model of relationship between apoptosis and autophagy in viruliferous whitefly and the consequence to survival rate and virus loading. TYLCV-induced apoptosis negatively regulated autophagy to maintain a homeostasis in favor of the coexistence of TYLCV within whitefly.

Considering canonical antagonism between apoptosis and autophagy and their contrasting effects on virus loading, the consequence of simultaneous stimulation in viruliferous whitefly for virus loading remain elusive. Our results showed that agonist-induced apoptosis negatively regulated autophagy, evidenced by accumulation of autophagy substrate SQSTEM1, and increased virus loading in whitefly. Conversely, inhibition of apoptosis reduced virus loading (Figure 6C, 6D). Once autophagy was agonist-induced or inhibitor-suppressed, however, virus loading was not affected by apoptosis but rather was significantly influenced autophagy status (Figure 6C, 6D). These results suggested that suppression of autophagy was required for the promotion of TYLCV loading in whitefly.

Stronger autophagy reduced virus abundance but was supposed to sacrifice the fitness of whitefly. On the contrary, stronger apoptosis favored virus loading within whitefly but could potentially be detrimental to whitefly. Assuming this was the case, PCD may function to maintain the intricate balance of whitefly fitness and virus loading, and our results experimentally tested this hypothesis (Figure 6E-G). By recording whitefly survival rate, we found that both suppression of autophagy by 3-MA and over-activation of autophagy by co-application of rapamycin and Z-VAD-FMK significantly reduced the survival rate of whitefly after 48 hours virus acquisition (Figure 6E). To rule out the effects of TYLCV infection, nonviruliferous whitefly was sampled to evaluate the influence of over-activation of autophagy on the survival rate of whitefly. By conducting a time-course records, we found decreased survival rate of nonviruliferous whitefly feeding on both rapamycin and Z-VAD-FMK for 36 h, suggesting that over-activation of autophagy can cause physiological costs in whitefly (Figure 6F). On the other side, overloading TYLCV by suppressing autophagy with 3-MA impaired the survival rate of whitefly (Figure 6G). Together, these findings suggested that both over-activated and suppressed autophagy were harmful to fitness of viruliferous whitefly. Only PCD with optimal balance of apoptosis and autophagy could maintain coexistence of TYLCV within whitefly without obviously physiological cost (Figure 6H).

## Discussion

PEBPs have been well characterized in its regulatory function related to host immunity, which involves multiple signaling pathways, such as MAPK, TBK, NFκB, and GSK (Gu et al., 2016; Sorriento et al., 2013; Su et al., 2016). The current study identified a novel PEBP in MED whitefly that served as a master regulator in coordination of apoptosis and autophagy *in vivo* to balance the virus loading and vector fitness. Genomics analyses reveals that whitefly genome consists of 202 PEBPs while 15 other arthropod species have 16 PEBPs at most (Chen et al., 2016). Further comparative transcriptome evidence shows that whitefly populations with stronger virus transmission had 20 PEBPs with higher expression than populations with weaker transmission ability, suggesting PEBPs are pivotal to virus transmission efficiency (Kliot et al., 2020). The complexity, however, has far-reaching consequences for our understanding of co-evolution of arbovirus and insect vectors. To our knowledge, this is the first study demonstrating that PCD immune homeostasis in shaping the balance between virus loading and vector fitness could be modulated by a single molecule from the insect vector, supporting the co-evolution theory of arbovirus with insect vector in nature.

### Regulation of PEBP-MAPK signaling cascade on apoptosis activation

The inhibition on Raf1 was firstly discovered on PEBP1, also called Raf1 kinase inhibitor protein (RKIP) (Yeung et al., 1999). Others PEBP, such as human PEBP4, can also suppress MAPK signaling cascade by interacting with Raf1 (Garcia et al., 2009; Hickox et al., 2002b). Our results showed that whitefly PEBP4, through interacting with Raf1 suppressed the phosphorylation of MAPK pathway. Therefore, a negative regulator of MAPK-apoptosis is also applicable for insect PEBPs. Furthermore, our results suggested that TYLCV CP stabilized PEBP4 and Raf1 interaction to suppress MEK/ERK phosphorylation. Since the phosphorylation status of PEBP could affect its regulatory function on multiple downstream signaling pathways, we propose that a complex of CP-PEBP4-Raf1 could be formed in viruliferous whitefly, by which CP may block the phosphorylate site of PEBP4 to prevent the dissociation of Raf1 from the complex (Corbit et al., 2003; Wen et al., 2014).

ERK, downstream of MAPK cascades, was typically characterized as transcriptional suppressor of pro-apoptotic factors and positive regulator of anti-apoptotic proteins, and high abundance of unbound pro-apoptotic factors usually leads to apoptosis (Kale et al., 2018; Lavoie et al., 2020). Our results demonstrated that silencing PEBP4 resulted in increased anti-apoptotic *Iap* and *Bcl-1* expression, indicating PEBP4 positively regulated apoptosis in whitefly. TYLCV CP augmented PEBP4 inhibition on MAPK pathway, and thereby reduced *Iap* and *Bcl-1* expression. Such negative transcriptional regulation explained TYLCV-activated apoptosis. In most cases, apoptosis in virus-infected hosts is induced by physical binding of extracellular signals to the transmembrane death receptor (Zhou et al., 2017). Alternatively, apoptosis could also be initiates via the mitochondria-dependent intrinsic pathway that mostly relies on the homeostasis between pro-apoptotic factors and anti-apoptotic factors (Bock and Tait, 2020). PEBP4-dependent activation of apoptosis in viruliferous whitefly apparently belonged to the latter, which broadly connected with other intracellular signals involved in differentiation, development, and homeostasis (Bock and Tait, 2020; Taylor et al., 2008).

Varied with the regulatory pathway involved, arbovirus species and vector tissue, apoptosis may have contrasting effects on arbovirus loading within insect vectors. The antiviral JNK pathway-activated apoptosis restricts the arboviruses Dengue, Zika and Chikungunya in salivary gland of *Aedes aegypti*, while current study shows that MAPK-dependent activation of apoptosis promoted TYLCV loading in MED whitefly (Chowdhury et al., 2020). Likewise, apoptosis occurred in midgut of *Ae. aegypti* increases Sindbis virus infection (Wang et al., 2012). Intriguingly, even in the brain of MED whitefly, a caspase-dependent apoptosis is induced by TYLCV and results in neurodegeneration, by which behaviorally promotes the TYLCV dissemination among host plants (Wang et al., 2020a). Considering the complex arbovirus-vector interactions, effects of apoptosis activation on arbovirus loading is subject of debate. The mechanistic role and unique pattern of apoptosis activation in insect vectors need to be illustrated because such information helps to understand how viruses survive from vector immunity.

### Direct and indirect regulation on autophagy

There are strong indications that PEBPs are involved in regulating autophagy-associated degradation (Noh et al., 2016; Zhao et al., 2020a). Human PEBP1 is reported to interact with LC3-interacting region (LIR) motif of PE-unconjugated LC3B, the homolog of ATG8 in mammals. The complex prevents the initiation of autophagy that is independent of MAPK signaling pathway (Noh et al., 2016). Moreover, human PEBP4 is co-localized with the lysosome, suggesting its potential role in regulation of degradation (Zhao et al., 2020a). Similarly, whitefly PEBP4 predictably consists of LIR motifs, capable of sequestering ATG8. Once hijacked by TYLCV CP, PEBP4 released ATG8 to initiate autophagy. Our competitive pull-down assays show that the intensity of PEPB4-dependent autophagy is enhanced with the increase of CP protein. By utilizing this pattern, unnecessary self-consumption may be largely avoided to prevent autoimmune diseases in insect vectors (Xu et al., 2020; Zhou and Zhang, 2012). Continuous feeding with virus-infected plant presumably exceeded the cell capacity for virions, possibly resulting in over-activation of autophagy and causing pathological lesions, and thereby increased whitefly mortality (Shintani and Klionsky, 2004). To deal with the damage of self-immunity, a negative regulation in equal intensity response to CP concentration, most notably through PEBP4-ERK-apoptosis signaling, is disclosed in viruliferous whitefly in ways that apoptosis counteracts the excessive activation of autophagy. It could provide a molecular explanation for arbovirus preservation within insect vector.

Different pre-domain sequences of PEBP4 from whitefly MEAM1 and from MED reasonably explained the results of Y2H and CoIP, as well as failure to screen any PEBPs in MEAM1 species that could interact with TYLCV CP (Zhao et al., 2020b; Zhao et al., 2020c). Although TYLCV could also induce PCD in MEAM1 species, the most striking difference is the initiation timepoint of PCD after acquiring TYLCV. Specifically, MED whitefly needs 6-12 hours to initiate PCD while MEAM1 needs a relative longer time from 24 to 48 hours (Wang et al., 2016; Wang et al., 2020c). Considering PCD could be essential for virus acquisition, PEBP4-dependent PCD in MED species has quicker initiation than PEBP4-independent PCD in MEAM1 species, possibly resulting in higher virus capacity for MED species (Kliot et al., 2020). This discovery will aid in understanding the rapid replacement of MEAM1 by MED species in Asia because arbovirus transmission could bring potential benefits for vector fitness (Eigenbrode et al., 2018).

### Immune homeostasis favors virus-vector coexistence

When challenged with pathogens, hosts are not only directly impaired by microbial invaders but usually cause cell and tissue damage because of overwhelming immunity (Martins et al., 2019; Oliveira et al., 2020). To prevent unnecessary costs of immune system, an alternative strategy widely adopted by plants to mammals is disease tolerance. It prefers to eliminate the deleterious effects produced by immunopathology rather than conferring direct immunity against infectious pathogens (Martins et al., 2019; Oliveira et al., 2020). In our study, a continuous activation of autophagy impaired the survival rate of whitefly even without TYLCV infection, revealing substantial physiological costs caused by immunopathology. By contrast, thoroughly immunosuppressed PCD by co-application of 3-MA and Z-VAD-FMK could amplify the burden of virulence, which subsequently impairs the fitness of vector (Oliveira et al., 2020). For instance, suppressing mosquito immunity leads to alphavirus over proliferation and increases the mortality of insect vector (Cirimotich et al., 2009; Myles et al., 2008). Similar with our results, inhibitor-suppressed autophagy in whitefly increased TYLCV loading but decreased survival rate of whitefly. Thus, a mild immunity is required for balancing arbovirus loading and vector fitness.

In conclusion, we have shown that TYLCV can simultaneously activate apoptosis and autophagy in MED whitefly via interacting with PEBP4. PEBP4 serves as a molecular switch to regulate both MAPK pathway relayed apoptosis and ATG8 initiated autophagy. The loading abundance of TYLCV within whitefly is dominantly determined by anti-viral autophagy because apoptosis that facilitating virus loading is arrested by autophagy. The homeostasis shaped by apoptosis and autophagy provides an immune tolerance, allowing TYLCV persistently preserved in whitefly population. These results provide an evolutionary insight into PCD regulations in vector that is more compatible with arbovirus. Based on these findings, more detailed understanding of molecular mechanism of immune homeostasis is of paramount importance for refining the utilization of immunoregulation approach for the future control of virus transmission.

## Materials and Methods

### Insect, plant, and virus

The infectious clone of *Tomato yellow leave curl virus* isolate SH2 (GenBank accession no: AM282874) was generously provided by Professor Xueping Zhou (State Key Laboratory for Biology of Plant Diseases and Insect Pests, Institute of Plant Protection, Chinese Academy of Agricultural Sciences). Tomato plants (*Solanum lycopersicum* cv Moneymaker), the natural host of TYLCV, were used for virus acquisition. Plants at the 3-4 true leaves stage were inoculated with the infectious clone, and confirmed by both visual observation and PCR analysis. All plants were reared at 26-28°C, with 60% relative humidity and a photoperiod of 16h : 8h (light/dark) cycle. The Mediterranean (MED) *Bemisia tabaci* (mtCOI GenBank accession no: GQ371165) were reared on cotton plants placed in insect-proof cages. New emerged adult whiteflies were randomly collected for the experiments.

### PCR and quantitative PCR (qPCR)

Total DNA of plants or insects was extracted using the RoomTemp™ Sample Lysis Kit (Vazyme, P073) according to manufacturer’s protocol. A 412 bp TYLCV fragment was PCR amplified using Taq PCR MasterMix (Tiangen, KT201) and primers V61 and C473 as previously described (Atzmon et al., 1998). Total RNA of whitefly samples was extracted by TRIzol™ Reagent (Ambion, 15596018), and reverse transcribed using the FastQuant RT Kit with gDNase (Tiangen, KR106). cDNAs of *PEBP4*, *PEBP4* pep1, *PEBP4* pep2, *V1* (TYLCV CP), *Raf1*, and *ATG8* were amplified by PrimerSTAR MAX DNA Polymerse (Takara, R045A). RT-qPCR reactions using the PowerUp™ SYBR Green Master Mix were carried out on the QuantStudio 12K Flex Real-Time PCR System (ABI) (ABI, A25742). Data were analyzed by the 2^−ΔΔCT^ relative quantification method, and each biological replicate consisted of three technical replicates.

### Yeast two-hybrid assays

The cDNA library of *B. tabaci* was constructed in plasmid pGADT7 (the prey), using the Clontech cDNA Library Construction kit (Takara, 630490) and transferred into yeast competent cells (strain Y187). The titer of the primary cDNA library was evaluated using the number of clones on plates, and the size of the inserted fragments in the library was confirmed by colony PCR. The full-length TYLCV V1 (CP) gene was cloned into PGBKT7 (the bait). The recombinant plasmid pGBKT7-CP was used to transform yeast strain Y2HGold and then selected on S.D./-Leu medium. Finally, the Y187 with prey plasmid and the Y2HGold with bait plasmid were screened with certain stress. Positive clones were selected on the triple dropout medium (TDO: S.D./-His/-Leu/-Trp) and quadruple dropout medium (QDO: S.D./-Ade/-His/-Leu/-Trp) with 3-aminotriazole (3-AT). All positive clones were sequenced and searched against the genome database of MED whitefly by Local Blastx 2.7.1 (Xie et al., 2017). Translated sequences were compared to the *Hemiptera* database in NCBI by Blastp. To verify the CP-PEBP4 interaction, full length PEBP4 was ligated into pGADT7 and co-transformed into yeast strain Y2HGold with pGBKT7-CP. Co-transformation of pGBKT7-lam and pGADT7-T vectors was used as a negative control. The presence of transgenes in yeast cells was confirmed by double dropout medium (DDO: SD/-Leu/-Trp). Triple and quadruple dropout media were used to assess the CP-PEBP4 interaction.

### Recombinant protein expression

To construct bacterial expression vectors, the amplified CD regions of *PEBP4*, *PEBP4* pep1 and *PEBP4* pep2 were cloned into pGEX-4T-1. The amplified CD regions of *V1* (TYLCV CP), *Raf1*, and *ATG8* were cloned into pET28a using *Xho* I and *EcoR* I sites. *Escherichia coli* BL21 (Takara, 9126) and BL21 (DE3) (Vazyme, C504) component cells were used for fusion protein expression. All constructs were verified by restriction enzyme digestion and DNA sequencing (BGI, China). Sequence of whitefly *PEBP4* and all primers mentioned above were list in *<u>Supplementary Table 2</u>*.

All recombinant plasmids (GST-PEBP4, GST-PEBP4 pep1, GST-PEBP4 pep2, His-CP, His-Raf1, His-ATG8) and empty plasmids (GST tag, 6×His tag) were transformed into component cells *Escherichia coli* BL21 (Takara, 9126) or BL21 (DE3) (Vazyme, C504), and induced by 0.1M IPTG for 4-16 hours at 16-37°C.

### Western blotting

Two rabbit anti-PEBP4 polyclone antibodies were prepared by BGI against synthetic peptides CPRKVRSRKNKENMES and TRHETTRSRPKNISPC. The commercial primary anti-Raf1 (D220484), anti-pMEK (D155070), anti-pERK (D151580), anti-SQSTM1 (8025S), anti-ATG8/GABARAP (13733S), and anti-GAPDH (60004-1-Ig) antibodies were respectively purchased from BBI, CST, and Proteintech. The production of rabbit anti-BtCaspase3b polyclone antibody has been described in our previous study (Wang et al., 2020a). The monoclonal antibody mouse anti-TYLCV CP was kindly provided by Professor Xiaowei Wang (Institute of Insect Sciences, Zhejiang University). HRP-conjugated GST tag monoclonal antibody (HRP-66001) and HRP-conjugated His tag monoclonal antibody (HRP-66005) were purchased from Proteintech. The commercial secondary antibodies goat anti-mouse IgG (ab6789) and goat anti-rabbit IgG (ab6721) were purchased from Abcam.

### GST pull-down assays

All GST pull-down assays were conducted by the GST Protein Interaction Pull-Down Kit (Pierce, 21516) according to the manufacturer’s protocol. In brief, after IPTG induction, total proteins extracted from recombinant expression cells were incubated with the agarose for 1-2h for GST-tagged proteins, and the His tagged fusion proteins were then incubated for 2-3h. In competitive interaction assays, His-ATG8 and His-CP were simultaneously incubated with GST-PEBP4-bound agarose. The glutathione agarose samples that hardly be eluted by 10-50mM glutathione was boiled within wash solution and SDS sampling buffer at 100°C, and discarded after centrifugation (mainly His-ATG8 and His-Raf1). The supernatant was finally analyzed by Western blotting. For boiled samples, nonspecific bands could appear in control groups due to the basal nonspecific interaction of input samples and glutathione agarose.

### Co-immunoprecipitation (Co-IP) and in vitro protein co-incubation assays

CoIP was conducted by the commercial CoIP Kit (Pierce, 26149) according to the manufacturer’s protocol. Total proteins from whiteflies were extracted in IP lysis buffer, and were subjected to the following assay. The final elution of CoIP was analyzed with Lane Marker Sample Buffer containing 100mM DTT in a reducing gel. For protein co-incubation assays *in vitro*, whitefly lysate was directly co-incubated with the fusion protein for 2-3 hours, and then, the precipitation was discarded after centrifugation. The final supernatant was analyzed by western blotting.

### Immunofluorescence detection and TUNEL assays

Collected whiteflies were fixed in 4% paraformaldehyde overnight and rinsed in TBST (TBS with 0.2% Triton X-100) for 30 minutes. For the TUNEL assay, samples were carried out by *in situ* Cell Death Detection Kit, POD (Roche, 11684817910) according to the manufacturer’s protocol. For immunofluorescence, samples were blocked with Pierce SuperBlock™ T20, and incubated with the primary antibody at room temperature for 3h. Following rinsing for five times in TBST (TBS with 0.05% Tween 20), samples were incubated with the secondary antibody at room temperature for 2 h. If two primary antibodies from the same host species were necessary, such as anti-Raf1, anti-PEBP4, and anti-ATG8, the sequential method was applied for multiple labeling in immunofluorescence. The anti-mouse IgG conjugate Alexa 488 (ab150113), anti-rabbit IgG conjugate Alexa 555 (ab150078), and anti-rabbit IgG F(ab’)_2_ fragment conjugate Alexa 594 (8889S) were purchased from Abcam and CST. Whitefly midguts and salivary glands were dissected after staining and mounted in the Fluoroshield Mounting Medium with DAPI (Abcam) before imaging under Zeiss LSM710 confocal microscope.

### Pharmacological experiments and RNA interference via feeding

Collected insects were placed in cylindrical light-proof containers with 30 mm diameter and 60 mm height. Each container was provided with 500 μl of 15% sucrose. MEK phosphorylation inhibitor mirdametinib (PD0325901) (Selleck, S1036), apoptosis inducer PAC-1 (Selleck, S2738), inhibitor Z-VAD-FMK (Selleck, S7023), and autophagy inducer rapamycin (Millipore, 553210) were dissolved in dimethyl sulfoxide (DMSO) and mixed in the diet at a concentration of 10μM. Autophagy inhibitor 3-MA (Selleck, S2767) was dissolved in double distilled water and supplied with the diet at 1μM. Equivalent DMSO was used as feeding control. All chemicals were administered via 24 h feeding.

The T7 RiboMAX Express RNAi System (Promega, P1700) was used to synthesize double-strand RNA according to manufacturer’s protocol. dsPEBP4 (50μl, 10 μg/μl) (dsGFP as control) was added to 450μl artificial diet to feed insects. The interference efficiency was measured by RT-qPCR. All insects in the same container were collected as an independent biological sample, and each treatment contained at least three replicates. All leaf-cages were applied on the same group (one infected combined one uninfected plant) for a replicate. To collect viruliferous whitefly, nonviruliferous whitefly was transferred to TYLCV-infected tomato plant for virus acquisition.

### Survival rate analysis

Whiteflies, 100 as a replicate, were fed in a feeding container for a 24 h, then transferred and caged on plant leaves for 48 h for TYLCV acquisition. Surviving whiteflies in each leaf-cage were recorded for survival rate calculation. In order to over-activate autophagy, a time-course feeding treatment was performed that lasted for 72 hours. At each time point, numbers of survivors in each of the three biological replicates were recorded to calculate an average survival rate. To verify if virus overloading would affect whitefly survival, autophagy was firstly suppressed by 3-MA for 24 h, and 60 individuals per biological replicate, were transferred and caged on TYLCV-infected vs. uninfected plants for another 48 h. Three replicates in total were carried out.

### Statistical Analysis

Statistical analyses were performed with SPSS software (Chicago, IL, USA). All data were checked for normality by the Wilk-Shapiro test. Two-tailed paired *t*-test was used to separate the means of normally distributed data, while Mann-Whitney test was used to analyze nonparametric distributed data. Asterisks denoted statistical significance between two groups (*P < 0.05, **P < 0.01, ***P < 0.001).

## Acknowledgments

We appreciate material support from Professor Xiaowei Wang and Jianxiang Wu, Zhejiang University, Professor Xueping Zhou, Institute of Plant Protection, CAAS, and Professor Chuanyou Li, Institute of Genetics and Development Biology, CAS. This project was supported by the National Key Research and Development Plan (2017YFD0200400) and the Strategic Priority Research Program of the Chinese Academy of Sciences (XDB11050400).

## Author contribution

Shifan Wang, designing project, performing experiments, analyzing data and writing manuscript; Huijuan Guo and Keyan Zhu-Salzman, manuscript writing; Feng Ge and Yucheng Sun, project designing and coordination, manuscript writing.

## Declare of interests

We declare no competing interest.

## Notes

### Competing Interest Statement

The authors have declared no competing interest.

