## Supplemental Information for "PEBP regulates balance between apoptosis and autophagy, enabling coexistence of arbovirus and insect vector"

**Supplementary Table 1**. Putative proteins of MED *Bemisia tabaci* that interacted with the coat protein of tomato yellow leaf curl virus in the yeast two-hybrid screening assay.

| Query No. | Score[a] | E-value[a] | Identity(%)[a] | Genome ID | Score[b] | E-value[b] | Identity(%)[b] | Acession No. | Description[c] |
|---|---|---|---|---|---|---|---|---|---|
| 1 | 25.0 | 5.4 | 43.3 | BTA008214.1 | 180 | 5.00E-47 | 27.72% | XP_018897595.1 | PREDICTED: uncharacterized protein LOC109030865 isoform X1 [Bemisia tabaci] |
| | | | | | 120 | 3.00E-27 | 24.76% | XP_018915628.1 | PREDICTED: glutamate receptor U1-like [Bemisia tabaci] |
| 2 | 94.4 | 4.80E-25 | 93.9 | BTA012003.1 | 263 | 9.00E-91 | 98.43% | XP_018901274.1 | PREDICTED: NADH dehydrogenase [ubiquinone] 1 beta subcomplex subunit 5, mitochondrial [Bemisia tabaci] |
| 3 | 149.0 | 1.82E-42 | 97.3 | BTA000233.1 | 516 | 0 | 96.73% | XP_018900154.1 | PREDICTED: uncharacterized protein LOC109032456 isoform X2 [Bemisia tabaci] |
| | | | | | 33.5 | 1.7 | 25.76% | RZF41284.1 | obscurin isoform X3 [Sipha flava] |
| 4 | 523.0 | 0 | 99.2 | BTA010352.1 | 1000 | 0 | 99.59% | XP_018911064.1 | PREDICTED: esterase E4-like [Bemisia tabaci] |
| 5 | 33.1 | 0.017 | 31.5 | BTA017005.1 | 564 | 0 | 96.50% | XP_018915181.1 | PREDICTED: probable dimethyladenosine transferase [Bemisia tabaci] |
| 6 | 26.2 | 6.5 | 71.4 | BTA006720.1 | 564 | 0 | 96.50% | XP_018915181.1 | PREDICTED: rho GTPase-activating protein 21 isoform X8 [Bemisia tabaci] |
| 7 | 265.0 | 2.89E-85 | 100.0 | BTA007379.1 | 567 | 0 | 99.62% | XP_018901717.1 | PREDICTED: alpha-N-acetylgalactosaminidase-like [Bemisia tabaci] |
| 8 | 227.0 | 4.41E-77 | 93.0 | BTA023875.1 | 267 | 4.00E-93 | 100.00% | XP_018910818.1 | PREDICTED: 40S ribosomal protein S15Aa-like [Bemisia tabaci] |
| 9 | 38.5 | 0.006 | 50.0 | BTA015065.1 | 286 | 4.00E-87 | 34.67% | XP_025417167.1 | ATP-binding cassette sub-family B member 10, mitochondrial [Sipha flava] |
| 10 | 157.0 | 2.10E-46 | 85.4 | BTA024002.1 | 293 | 2.00E-102 | 93.79% | XP_018899372.1 | PREDICTED: WAS/WASL-interacting protein family member 1-like [Bemisia tabaci] |
| 11 | 311.0 | 2.15E-97 | 99.3 | BTA011564.1 | 1976 | 0 | 99.36% | XP_018904225.1 | PREDICTED: C-1-tetrahydrofolate synthase, cytoplasmic [Bemisia tabaci] |
| 12 | 223.0 | 6.24E-69 | 94.1 | BTA012527.1 | 913 | 0 | 84.84% | XP_018917484.1 | PREDICTED: uncharacterized protein LOC109044293 [Bemisia tabaci] |
| | | | | | 34.7 | 1.9 | 27.63% | XP_027848922.1 | ethylmalonyl-CoA decarboxylase-like isoform X1 [Aphis gossypii] |
| 13 | 28.9 | 3.5 | 44.4 | BTA029690.1 | 2754 | 0 | 99.85% | XP_018903654.1 | PREDICTED: uncharacterized protein LOC109034784 [Bemisia tabaci] |
| | | | | | 1675 | 0 | 98.90% | XP_018903657.1 | PREDICTED: PAX3- and PAX7-binding protein 1 [Bemisia tabaci] |
| 14 | 106.0 | 2.57E-27 | 99.0 | BTA014874.1 | 1469 | 0 | 98.28% | XP_018901035.1 | PREDICTED: proteoglycan 4-like [Bemisia tabaci] |
| 15 | 145.0 | 1.14E-43 | 97.5 | BTA017133.1 | 518 | 0 | 99.21% | XP_018902549.1 | PREDICTED: probable 39S ribosomal protein L24, mitochondrial [Bemisia tabaci] |
| **16** | **306.0** | **1.08E-104** | **99.3** | **BTA013106.1** | **722** | **0** | **97.51%** | **XP_018910109.1** | **PREDICTED: uncharacterized protein LOC109039182 isoform X1 [Bemisia tabaci]** |
| | | | | | **336** | **3.00E-116** | **98.08%** | **XP_018910114.1** | **PREDICTED: phosphatidylethanolamine-binding protein homolog F40A3.3-like isoform X4 [Bemisia tabaci]** |
| 17 | 26.9 | 0.43 | 44.4 | BTA020855.1 | 459 | 9.00E-166 | 99.55% | XP_018897453.1 | PREDICTED: uncharacterized protein LOC109030777 [Bemisia tabaci] |
| | | | | | 126 | 1.00E-34 | 39.41% | VVC29685.1 | Immunoglobulin subtype,Immunoglobulin-like domain,Immunoglobulin-like fold [Cinara cedri] |
| 18 | 62.4 | 2.41E-10 | 34.4 | BTA027805.1 | 347 | 3.00E-108 | 41.13% | QEP09172.1 | cytochrome c oxidase subunit I [Aphaena discolor nigrotibiata] |
| 19 | 27.3 | 0.53 | 39.3 | BTA004184.1 | 4908 | 0 | 98.87% | XP_018899018.1 | PREDICTED: uncharacterized protein LOC109031748 [Bemisia tabaci] |
| | | | | | 957 | 3.00E-173 | 65.36% | XP_025414375.1 | protein FAM135A [Sipha flava] |
| 20 | 26.2 | 1.7 | 41.9 | BTA000356.1 | 1333 | 0 | 98.76% | XP_018901482.1 | PREDICTED: glucose dehydrogenase [FAD, quinone]-like [Bemisia tabaci] |
| 21 | 25.4 | 1.8 | 39.1 | BTA026158.1 | 623 | 0 | 99.35% | XP_018915405.1 | PREDICTED: facilitated trehalose transporter Tret1-like isoform X1 [Bemisia tabaci] |
| 22 | 29.6 | 4 | 34.5 | BTA021511.2 | 1143 | 0 | 96.05% | XP_018915012.1 | PREDICTED: cytochrome P450 4g15-like isoform X1 [Bemisia tabaci] |
| 23 | 27.3 | 0.41 | 41.9 | BTA019316.1 | 1060 | 0 | 100.00% | XP_018913374.1 | PREDICTED: kelch-like protein 30 [Bemisia tabaci] |

a Query sequences were used local Blastx to identify full amino acid sequences in MED whitefly genome data.

b Selected proteins were used Blastp and compared with the database of *Hemiptera* to predict putative function.

c If the subject protein was predicted as "uncharacterized protein", a further PSI-Blast result with specific description would be supplied below the previous result.

**A**

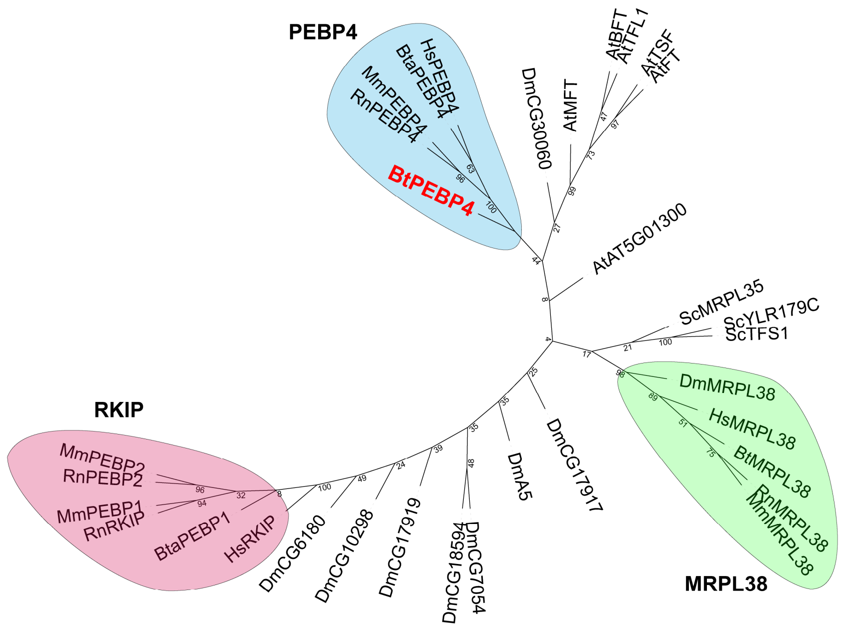

**B**

```
BtPEBP4    MASVSVMLSMVLQTYSVSVVSMAHSLHLLGFPPRKVRSRKNKENMERTDSPESMEMESTKGTESTKKTRRTKSTKTKSTK   80

BtPEBP4    SKKPKSTKSTKTKSTKARSTKITKTKKTPTPKIKKSTRNTRRPRISKNTESTKISKTTEVIGSTEILTTTRHGKTRSRPK  160
HsPEBP4    ......................................MGWTMRLVTAALLLGLMMVVTGDEDE...............N   27
MmPEBP4    ...............................MTMKLVAAALCLSLLAAGLWVGLSLTAESIEEGKPGGEKPGGGKPGGSG   49
BtaPEBP4   ......................................MGWTMRLVTAALLLGLAVAVTGEEED...............A   27
RnPEBP4    .................................MKLVAAAFGLSLLVAGLWVGLSLTSNSDGGGKPGG...GGG..GGGG   42

BtPEBP4    NISPFDAIFRENEIVPQIVNNTPIYKCEVFYGKEKIHFGNNMTADQIAEKPTLVKWPTQSG.AFYTLLMFGPDVPTREKS  239
HsPEBP4    SPCAHEALLDED..TLFCQGLEVFYPELGNIGCKVVPDCNNYRQKITSWMEPIVKFPGAVDGATYILVMVDPDAPSRAEP  105
MmPEBP4    RGCFLPPLPKED..VSLCRNLEVFYMEMGNISCKIVPKCNLYRQKITAWQAPIVKFHTALDGALYLLVMVDPDAPSRSNP  127
BtaPEBP4   DLCVYEALPDND..AVLCKGLKVFYPELGNIGCMIVPECNNYRQKITTWPEPIVKFPQALDDAAYILVMVDPDAPSRSSP  105
RnPEBP4    RGCFLPPLPKED..VSLCRNLEVFYMEMGNISCKVVPKCNQYRRKIMTWQSPIVKFHGALD...................  101

BtPEBP4    CEMMWRHWAVGNIPKND....IHKGDTLVEYENIEPFYRAGFHRILFFIYKQPGRRRLNFTSSKGVIKKGAKWEVRGVWY  315
HsPEBP4    RQRFWRHWLVTDIKGADLKKGKIQGQELSAYQAPSPPAHSGFHRYQFFVYLQEGKV.ISLLPKENKTRG........SWK  176
MmPEBP4    VMKYWRHWLVSNITGADMKSGSIRGNVLSDYSPPTPPPETGLHRYQFFVYLQGDRD.ISLSVEEKADLG........GWN  198
BtaPEBP4   KARFWRHWLVSDIKGSDMKIGKIQGQELSPYHPPSPPAHSGLHRYQFFVYLQEGRT.ISLSPKENKTRG........SWK  176
RnPEBP4    .........VSELC.......WLKENVGPQLAFSSPGP..........VFLLTN...IKWPAFENVSL............  140

BtPEBP4    IERFLAKFNLTEPIAANFFFLYDLNPHPEIPQVTVDPLSLIR.........                              357
HsPEBP4    MDRFLNRFHLGEPEASTQFMTQNYQDSPTLQAPRERASEPKHKNQAEIAAC                              227
MmPEBP4    LDKFLQQYGLRDPDTSTQFMTQFDEELSSEFGRINDDQEQFNQK.......                              242
BtaPEBP4   MDKFLSHFHLTEPEASTQFMTQYYLDGLSRQNPGVVSNETTDKPEPK....                              223
RnPEBP4    ...FSSHRKLKSPKERLD.................................                              155
```

**Supplementary Figure 1. Homology analysis of the candidate whitefly PEBP.** (A) The PEBP protein sequences that identified from 7 species and the candidate PEBP in whitefly were taken as input sequences, and the phylogenetic tree was constructed through the neighbor-joining (N-J) method with 1000 bootstrap replicates. Three major clusters were named according to the dominate proteins. (B) The candidate PEBP in whitefly was named as BtPEBP4, and the amino acid sequence was aligented with other PEBP4s in mammals.

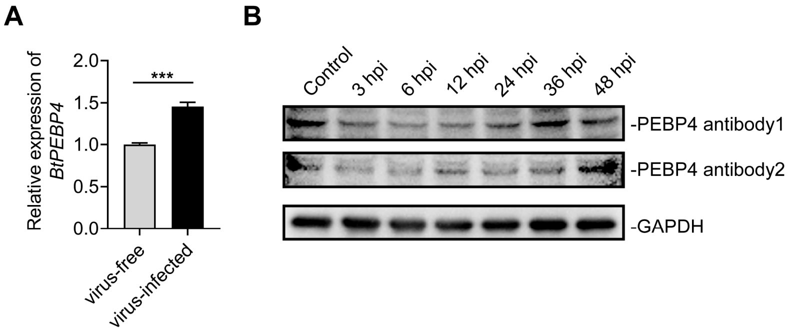

**Supplementary Figure 2. TYLCV enhances the relative abundance of PEBP4 in whitefly.** (A) After 48 hours TYLCV acquisition, the transcriptional expression of *BtPEBP4* was analyzed by qPCR, n=5. (B) The dynamic of the PEBP4 protein abundance in whitefly after virus acquisition was analyzed by immunoblotting.

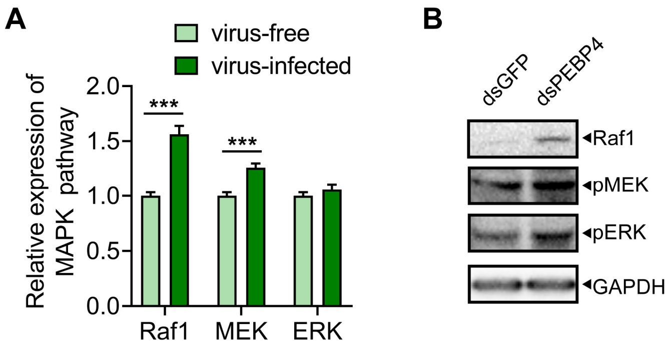

**Supplementary Figure 3. Both TYLCV and PEBP4 mediates MAPK pathway.** (A) The transcriptional expression of *Raf1*, *MEK*, and *ERK* after 48 hours TYLCV acquisition, n=5. (B) The phosphorylation of MAPK pathway after PEPBP4 knock-down was analyzed by immunoblotting.

iLIR Autophagy Database

Query: >PEBP4

| Motif | Start | End | Pattern | PSSM Score | LIR in Anchor |
|---|---|---|---|---|---|
| WxxL | 163 | 168 | SPFDAI | 9 | Yes |
| WxxL | 221 | 226 | GAFYTL | 5 | No |
| WxxL | 264 | 269 | VEYENI | 9 | No |
| WxxL | 275 | 280 | AGFHRI | 6 | No |
| WxxL | 331 | 336 | ANFFFL | 7 | No |

**ANCHOR Positions - Start and End**

| Anchor | Start | End |
|---|---|---|
| 1 | 1 | 38 |
| 2 | 96 | 103 |
| 3 | 137 | 148 |
| 4 | 162 | 175 |
| 5 | 179 | 189 |

**Supplementary Figure 3. Putative LIR motif of PEBP4.** LIR motifs are necessary for the direct interaction with autophagy related ATG8 family. The amino acid sequence of whitefly PEBP4 was analyzed and predicted by the online tool of LIR motif searching in iLIR Autophagy Database (https://ilir.warwick.ac.uk/index.php).

**Supplementary Table 2. Primer List.**

| Name | Primer Sequence (5'-3') | Accession No. |
|---|---|---|
| *Actin* | TCTTCCAGCCATCCTTCTTG | XM_019042718.1 |
| | CGGTGATTTCCTTCTGCATT | |
| *Caspase1* | TGTTGGAGACGGTATGGA | BTA009205.1 |
| | ATGAAGACAGTGCTTAATGC | |
| *Caspase3* | CATCACGATCAACGGGACCA | BTA015946.1 |
| | TGTCGATGTGCTGCTCGAAT | |
| *TYLCV* | GAAGCGACCAGGCGATATAA | AM282874.1 |
| | GGAACATCAGGGCTTCGATA | |
| *PEBP4* | CCAGGAAGAAGGAGATTGAA | BTA013106.1 (Sequenced PEBP4 CDs region is appended below) |
| | GCTATTGGTTCGGTCAGAT | |
| *Iap* | ATGTCTCGGAATCATCTCAG | BTA000132.1 |
| | ACCACAAGGAAGGAATGC | |
| *Bcl-2* | GCTAATGACACAGACTGGAT | BTA025366.1 |
| | GAGATGAAGTTCGTGAGGAA | |
| *Raf1* | AAGTGCTGATGATAGTGCTA | BTA014695.1 |
| | CATCGGTAGACAGTTCCAA | |
| *MEK* | TTGAGTTGCTGGACTACATAG | BTA026985.1 |
| | TTATCCGCCTCGCACTTA | |
| *ERK* | CAATGATAAGGCACGCAAT | BTA017617.1 |
| | CATAATACTGTTCCAGGTAAGG | |
| *ATG3* | CGTTTAAGGGAACAGCACTTG | BTA008054.1 |
| | CCAGATTGTCTCCAGCAGCA | |
| *ATG9* | TTGCCATCATTAACTTTCTGCT | BTA017980.1 |
| | AGGGTTCCTGGTTCACGC | |
| *ATG8* | TACACTTGAGACCAGAGGA | BTA002927.2 |
| | CTTCTTCGTGATGTTCTTGA | |
| *ATG12* | TCAAAGCCACTGGTAACGC | BTA009484.1 |
| | TCTGGTCCGGAGCAGGAGC | |
| dsGFP | CTCGTGACCACCCTGACCTAC | |
| | GTTCACCTTGATGCCGTTCTT | |
| | T7-CTCGTGACCACCCTGACCTAC | |
| | T7-GTTCACCTTGATGCCGTTCTT | |
| dsPEBP4 | CAACCACAAGGCACGAA | |
| | TCCTTCTTCCTGGCTGTT | |
| | T7-CAACCACAAGGCACGAA | |
| | T7-TCCTTCTTCCTGGCTGTT | |

| | | |
|---|---|---|
| PEBP4 | CGCGAATTCATGGCTTCCGTCTCCGTAAT | EcoRI/XhoI |
| (BL21) | CGCCTCGAGTCATCGAATTAGTGACAGTGG | (pGEX-4T-1) |
| PEBP4 pep1 | CGCGAATTCATGGCTTCCGTCTCCGTAAT | EcoRI/XhoI |
| (BL21) | CGCTCGAGTCAAGGCCACTTGACTAACGTTG | (pGEX-4T-1) |
| PEBP4 pep2 | GCGAATTCACGCAGAGTGGTGCTTTTTA | EcoRI/XhoI |
| (BL21) | CGCCTCGAGTCATCGAATTAGTGACAGTGG | (pGEX-4T-1) |
| Raf1 | CGCGGAATTCATGTCTGTCGAATATGACGA | EcoRI/XhoI |
| (BL21(DE3)) | CGCCTCGAGTTAGATTATTCCACCCATTG | (pET28a) |
| ATG8 | CGCGCGGAATTCATGAATTTCCAATACAAAGC | EcoRI/XhoI |
| (BL21(DE3)) | CGCTCGAGTTAAGCACAGATCTGGC | (pET28a) |

T7=5’-TAATACGACTCACTATAGG-3’

>BtPEBP4

ATGGCTTCCGTCTCCGTAATGCTATCCATGGTTTTGCAGACGTATTCGGTGTCCGTGGTATCTATGGCACATTCGTTGCATTTGCTCGGTTTTCCCCCGAGGAAAGTGAGAAGCAGGAAAAACAAGGAAAACATGGAAAGAACGGATAGCCCGGAAAGTATGGAGATGGAAAGCACGAAAGGCACAGAAAGCACAAAGAAAACAAGACGCACGAAAAGCACAAAAACTAAAAGCACGAAAAGCAAAAAACCCAAAAGCACGAAAAGCACAAAAACCAAAAGCACAAAAGCCAGAAGCACGAAAATCACAAAAACAAAAAAAACACCAACACCAAAAATCAAGAAAAGCACAAGGAACACGAGGCGTCCGCGAATAAGTAAAAACACGGAAAGCACCAAAATTTCAAAAACCACTGAAGTTATTGGAAGCACAGAAATTTTAACAACCACAAGGCACGGAAAGACGAGGTCAAGACCTAAAAATATATCACCTTTTGATGCAATTTTTAGAGAAAATGAAATAGTGCCACAGATCGTTAATAACACACCAATATACAAATGTGAGGTTGAATATGGCAAGGAGAAAATACATTTTGGGAACAACATGACGGCAGATCAAATCGCAGAAAAACCAACGTTAGTCAAGTGGCCTACGCAGAGTGGTGCTTTTTATACTCTTCTCATGTTTGGACCTGATGTCCCAACGCGGGAGAAATCTTGTGAGATGATGTGGCGACATTGGGCAGTTGGGAATATCCCAAAGAATGACATCCATAAAGGAGACACCTTAGTCGAATACGAAAATATCGAGCCTTTCTACAGGGCAGGTTTTCACAGGATATTGTTTTTTATTTACAAACAGCCAGGAAGAAGGAGATTGAACTTCACATCAAGTAAAGGGGTTATTAAGAAAGGAGCGAAATGGGAAGTGCGAGGAGTCTGGTATATTGAGCGTTTTCTCGCTAAATTCAATCTGACCGAACCAATAGCAGCGAACTTTTTCTTTCTTTATGATTTGAATCCTCATCCCGAAATTCCACAAGTTACTGTAGATCCACTGTCACTAATTCGATGA
